# Spatial proteomics reveals heterogeneity in neural markers underpinning high-fat diet-induced myopathy in male mice

**DOI:** 10.1101/2023.06.12.544330

**Authors:** Lydia Hardowar, Jayakumar Vadakekolathu, Sergio Rutella, Richard P. Hulse, Craig L. Doig

## Abstract

Metabolic dysfunction in skeletal muscle disturbs its contractile response as well as its innervation and vascular networks. The molecular drivers responsible for affecting decline in function remain poorly defined. To provide insight and locate these, we mapped changes in the spatial proteome occurring as a result of impaired metabolic health. We exposed male mice (C57/BL6J) to diet induced obesity to investigate the impairment of muscle metabolic function and myopathy. We conducted digital spatial profiling using the NanoString GeoMx® platform on recovered skeletal muscle (*tibialis anterior*) comparing it to standard fed controls. Digital spatial profiling revealed areas with shifts in the contractile protein desmin and CD31 expression, a marker of tissue stress and cellular maladaptation. We find increased expression of proinflammatory markers were identified in areas of elevated Desmin in obese samples compared to controls. Our data suggest a dietary-driven relationship between the spatial abundance of the sarcomere protein desmin and the influx of neural and inflammatory mediators to muscle. This supports the concept of pro-inflammatory events underpinning the muscle metabolic dysfunction associated with chronic non-communicative diseases such as type 2 diabetes, metabolic syndrome, and chronic obstructive pulmonary disorder.

## Introduction

Sustained accumulation of body mass contributes to a range of metabolic dysfunctions. Increased risks of cardiovascular disease, cancer and widespread metabolic failure emphasise the importance diet plays in tissue health. Amongst obesity-related comorbidities, myopathy is a critical element affecting the global population (McCormick and Vasilaki, 2018; Morgan and Partridge, 2020; Nathan and Fuld, 2010).

The burden of increased lipid availability to skeletal muscle and subsequent shifts in energetic demand have profound impacts upon a tissue’s ability to preserve architecture and sustain homeostasis. Physiological decline associated with increased body fat mass is thought to be partially driven by infiltrating mediators alongside parallel degradation of tissue-tissue interfaces. Examples in skeletal muscle, comprise of immune cell invasion and loss of neuromuscular junctions. The neuro-inflammatory events associated with muscle dysfunction are shown to occur alongside widespread functional decline (Christ and Latz, 2019; Fink *et al*., 2014; Heinrich *et al*., 2015). However, the molecular understanding of these events is obfuscated by tissue-tissue interactions. This represents a barrier to the development of interventions to preserve skeletal muscle function and improved understanding of obesity related disease. (Samdani *et al*., 2015).

Desmin is a key myofibrillar protein responsible for regulation of sarcomere architecture. Expression of desmin is not uniform through muscle tissue with enrichment at nerve-muscle borders including neuromuscular junctions (NMJs) (Elashry *et al*., 2019; Mogi *et al*., 2016) (Eiber *et al*., 2020). Regional changes in desmin abundance have been recorded and are linked to shifts in mechanical force and diets high in fat reduce desmin levels (Palmisano *et al*., 2015). Desmin abundance in human skeletal muscle is also significantly reduced in those living with type 2 diabetes (Hwang *et al*., 2010). Using high fat diet to induce obesity in mice damages NMJs (Martinez-Pena y Valenzuela and Akaaboune, 2020a). Despite this desmin impacts in the response to high-fat diet are unclear, as are any changes in its spatial abundance.

Angiogenic and microvasculature performance are also both fundamental to skeletal muscle performance. Endothelial cells form structures to preserve blood flow and their dysfunction is associated with diet-induced myopathy. Diets containing excess caloric content originating from fat have been shown to increase the abundance of the endothelial marker CD31 (Heinrich *et al*., 2015). This suggests shifts in protein abundance found within CD31 containing muscle are integral to the development of pathogenic states. We set out to address this by inducing metabolic syndrome in a commonly utilised murine model. We then collected skeletal muscle tissue that was subjected to spatial cell profiling using digital proteomics. By using a targeted protein panel we provided insight into the main neuronal and inflammatory changes associated with obesity related muscle wasting. This work reveals spatial attributes of vascular and muscular tissue mediated through lifestyle behaviours. It also identifies specific morphology key to metabolic syndrome development.

## Methods and Materials

### Animals

Mice (C57/BL6J) were purchased from Charles Rivers, UK. They were group housed in humidity and temperature (22°C) controlled conditions with a 12:12 light/dark cycle. All mice within study were purchased age-matched at 6 weeks old. Nesting material was provided in the cages along with enrichment in the form of chew sticks and ad libitum access to water and food. All mice were maintained, and tissue samples were taken in accordance with protocols approved by the local AWERBs at University of Nottingham and Nottingham Trent University and carried out under the Animals (Scientific Procedures) Act 1986.

### Inducing hyperglycaemia with a high fat diet

Six male C57/Bl6J mice were purchased from Charles River Laboratory. Animals were provided with Rodent 2018 Envigo Global Certified Diet (n=3) (Envigo Laboratories U.K. Ltd., Oxon, UK; 6.2% by kcal of fat content) standard chow or 60% by kcal high fat diet (n=3) (Teklad custom diet TD.06414; Envigo Laboratories U.K. Ltd., Oxon, UK) for the experimental diet for 12 weeks.

### Tissue processing

Mice were culled by schedule 1 cervical dislocation. *Tibialis anterior* skeletal muscle tissues were removed from animals and cryoprotected in 30% sucrose overnight. Tissues were embedded in optimal cutting temperature compound and stored at -80°C. Tissue samples were cut serially sectioned at 30μm using the Leica cryostat (CM1860 UV) and placed across 10 SuperFrost microscope slides (VWR International; UK).

### Immunofluorescent staining

Tissue sections were thawed to room temperature and washed in 0.01M phosphate buffer saline, permeabilised in 0.2% Triton and blocked with 5% bovine serum albumin at room temperature for 1 hour. Following from this, primary antibodies of 1:200 Goat anti-CD31 (R&D systems, U.K, goat anti-PECAM1) and 1:500 anti-desmin (Abcam, U.K, Alexa Fluor™ 647 anti-desmin, ab195177) diluted in 5% BSA were incubated on tissue for 72 hour at 4°C. Following this, tissues were washing thrice with PBS and incubated with Alexa Fluor™ 488 Donkey anti-Goat IgG (ThermoFisher Scientific, U.K) for 2 hours and washed again with a final DAPI staining. Coverslips were mounted onto slides and imaged using the Leica SP5 confocal. The oil emersion x20 magnification objective was used to image the skeletal tissue sections. X20 magnification was used as representative images. 20 frames per images within a Z-stack image were taken. Images were transported and processed within Fiji software for further analysis.

### Confocal image analysis

Confocal captured images were processed within ImageJ v. 2.0.0. software. Images were opened as Z-stack hyperstacks. All Images to be analysed were calibrated with a reference image including a scale bar. Stacks with no fluorescent signalling were removed and the rest of the hyperstack were converted into a Z projection to form a 2-dimensional image. The contract, background, minimal and maximal signal on a 0-250 grey value scale were adjusted in each signal channel and kept consistent between each image in each treatment group analysed. Within ImageJ, integrated density of the CD31 and desmin separated channels were analysed by drawing a square from the selection tool around 1024×1024 pixel images. The integrated density measurement setting was applied and values were extracted from results table. Background integrated values were subtracted from each image integrated density. For muscle fiber cross-sectional area measurement units were displayed as μm^2^.

### GeoMx® Digital Spatial Profiling of *Tibialis anterior* skeletal muscle tissue

GeoMx® Digital Spatial Profiling (DSP) technology (Nanostring; Seattle WA) was utilised to quantify a protein array of *tibialis anterior* muscle skeletal muscle tissue from standard chow and 60% high fat diet fed mice.

Following on from the tissue processing steps, tissue sections (5μM thickness) were washed in TBST for 10 minutes three times. Tissue sections were incubated overnight at 4°C with a cocktail diluted in Buffer W (Nanostring; Seattle WA) containing morphology markers Desmin (Abcam, U.K, Alexa Fluor® 647 anti-desmin, ab195177) and CD31 (R&D systems, U.K, goat anti-PECAM1) and UV-cleavable oligo-labelled detection antibody panel (GeoMx Neural Cell Profiling Panel). Tissues were washed with TBST and incubated with secondary antibody Alexa Fluor 488 mouse anti-goat for 2 hours. Following this, tissues were washed with TBST and incubated for an additional 15 minutes with nuclear stain SYTO 83 (1/10,000). A final wash was given with 1X TBST and tissues were transferred to GeoMx**®** DSP instrument. Once scanned by the instrument, morphological markers were used to select for geometric segment regions of interest (ROIs). Geometric segment ROIs included 400μm diameter circular selection of high desmin and high CD31 regions of all tissue sections per animal across both experimental groups. DSP UV-illuminator cleaved and released the conjugated oligonucleotide-labels from the geometric segment ROIs selected. The oligo-labels from each ROI were counted using nCounter analysis system using a high-resolution scan setting (555 FoV).

### Data processing and analysis

The count data (RCC files) generated using the nCounter digital analyser transferred into the GeoMX DSP analyser for further quality controls and analysis. The data was normalised to the internal controls of each ROI (ribosomal protein S6, histone H3 and GAPDH) by GeoMx**®** DSP software (v2.4). Differential expression plotted as Log2 transformed and normalized counts.

### Statistical analysis

All data were analysed in GraphPad Prism Version 9 and reported as the mean ± standard error for mean (S.E.M). Experimental group differences from the image analysis data of Desmin and CD31 immunohistochemistry were statistically determined by unpaired t-test for comparisons of integrated density and cross-sectional area measurements. Data from geometric segment ROI counts were separated by experimental groups (standard chow vs. high fat diet) were analysed by one-way ANOVA.

## Results

### Verification of high fat diet impacts on skeletal muscle

Mice were fed either a high fat diet or a normal chow diet for 12 weeks. Mice fed high fat diet accumulated weight and exhibited elevated blood glucose levels by week 12 in comparison to normal chow (Fig 1A-B). High fat diet impacts over gross muscle fibre architecture were verified by a demonstration of reduced *tibialis anterior* cross-sectional area (CSA), determined by stained muscle fibers (Fig. 1C). This is consistent with documented impacts of high fat diet and elevated lipid exposure reducing the CSA (Elashry *et al*., 2019).

**Figure 1.**
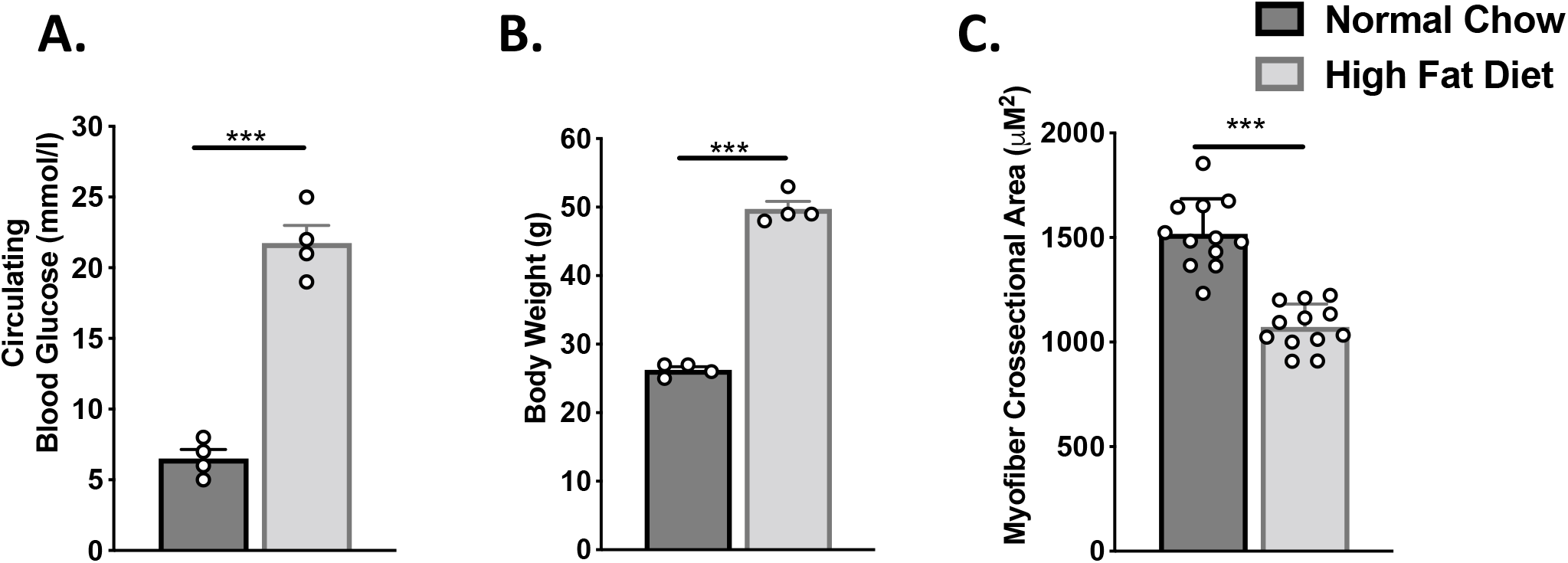
Establishment of high fat diet induced myopathy. C57/BL6J mice (6 weeks old) were randomly assigned to groups fed normal mouse chow (n=3) or high fat diet (n=3) for 12 weeks. (A) Circulating blood glucose levels from mice was recorded at 12 weeks. (B) Total body weights from mice at 12 weeks. (C) Skeletal muscle fiber circumference of tibialis anterior muscle from high fat diet and normal diet mice. Data presented as means ±SEM. *P<0.05, **P<0.01, ***P<0.0001.

### Digital spatial profiling of high fat diet and normal chow skeletal muscle

*Tibialis anterior* muscle was collected from these mice and subject to spatial proteomics described in the schematic workflow (Fig. 2A). Targeted quantification of a panel of neural markers was used to measure proteins responsible for the major driving events of a loss of contractile function. Digital protein expression data were recovered from muscle (Fig. 2B), this was converted to log2 scaled data (Fig. 2C). Abundance of the endothelial protein CD31 was shown to be significantly elevated in muscle of high fat diet mice, compared to normal chow. Similarly, the microtubule associated protein 2 (MAP2) neurofilament protein is also induced by high fat. Using the quantitative analysis of the DSP we also detected significantly increased changes across tissue in the abundance of neuronal cell markers CD68, NeuN, S100B, TMEM119 and Olig2. The proliferation mark Ki-67 was also increased. These data establish spatially resolved changes in the abundance of neuronal proteins as a response to a high fat diet.

**Figure 2.**
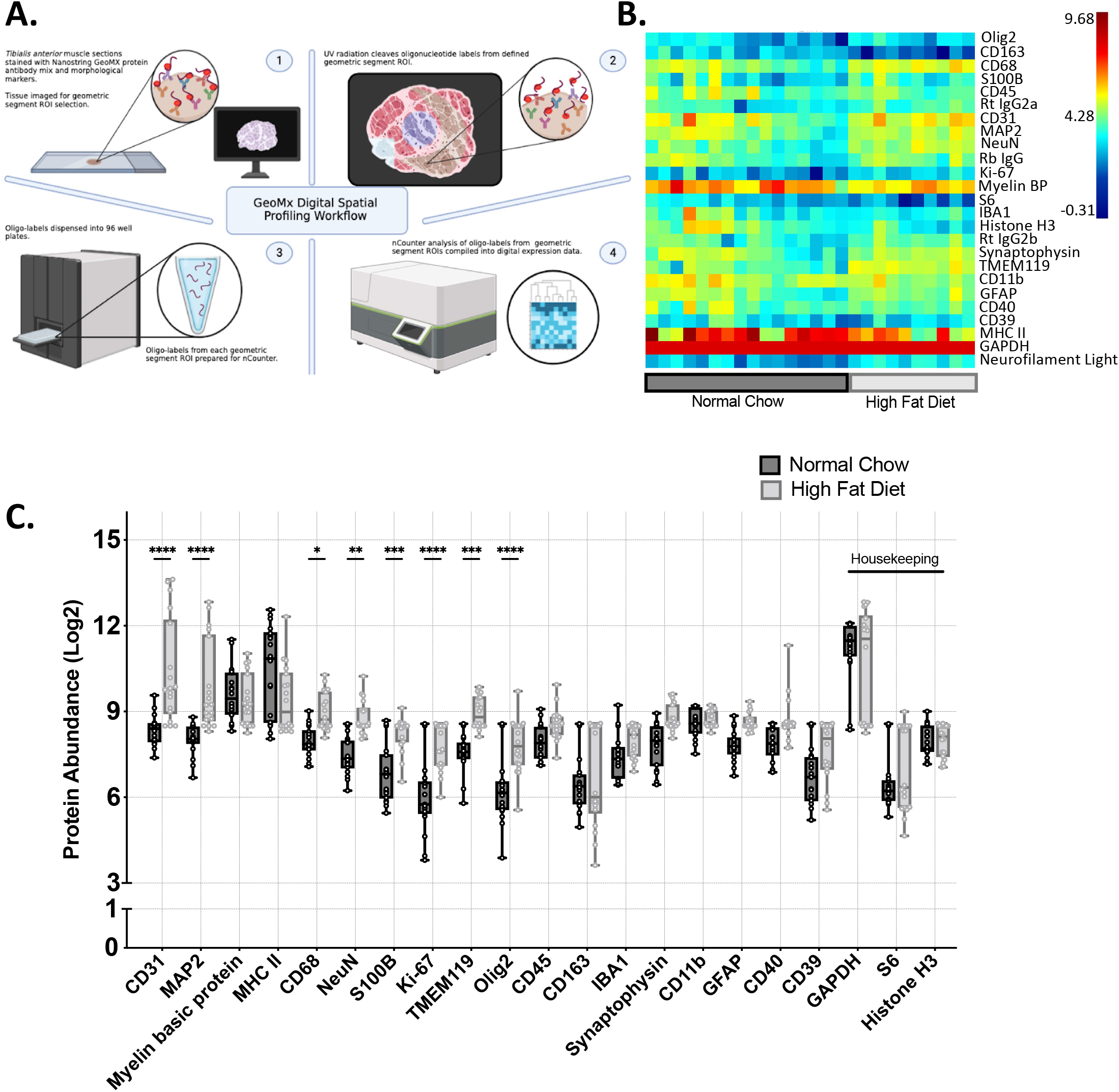
Digital Spatial proteomics of high fat diet induced myopathy in mice. (A) Schematic of the digital spatial proteomics workflow. (B) Heatmap of the protein abundance data from the GeoMX DSP. (C) Protein abundance (log2) of targeted panel in muscle of mice fed a normal or high fat diet. Data presented as box and whisker plots (n=10-16) ±SEM *P<0.05, **P<0.01, ***P<0.0001.

### Skeletal muscle tissue exhibits spatial heterogeneity in response to high fat diet

We validated and reproduced the DSP outcomes through immunological staining for CD31 in normal and high fat diet fed muscle tissue (Fig. 3A-B). Counts for CD31 were significantly higher in the high fat diet mice when assessed by digital spatial proteomics and immunostaining using conventional antibody generated data (Fig. 3C). This agrees with DSP obtained data and is supportive of a CD31 mediated response to the high fat diet induced changes in muscle architecture and endothelial infiltration during metabolic stress (Pi *et al*., 2018). Cell nuclei were also stained for using SYTO83 and desmin was used as an indicator of myofiber structure. Intensity of desmin staining was reduced in these mice compared to normal chow (Fig. 3D). Using *in situ* spatial proteomics we identify these specific regions of interest with desmin enrichment or CD31 depletion. To document and quantify these changes in muscle and endothelial markers we divided loci dependent upon their abundance of CD31 or desmin (Fig. 3E).

**Figure 3.**
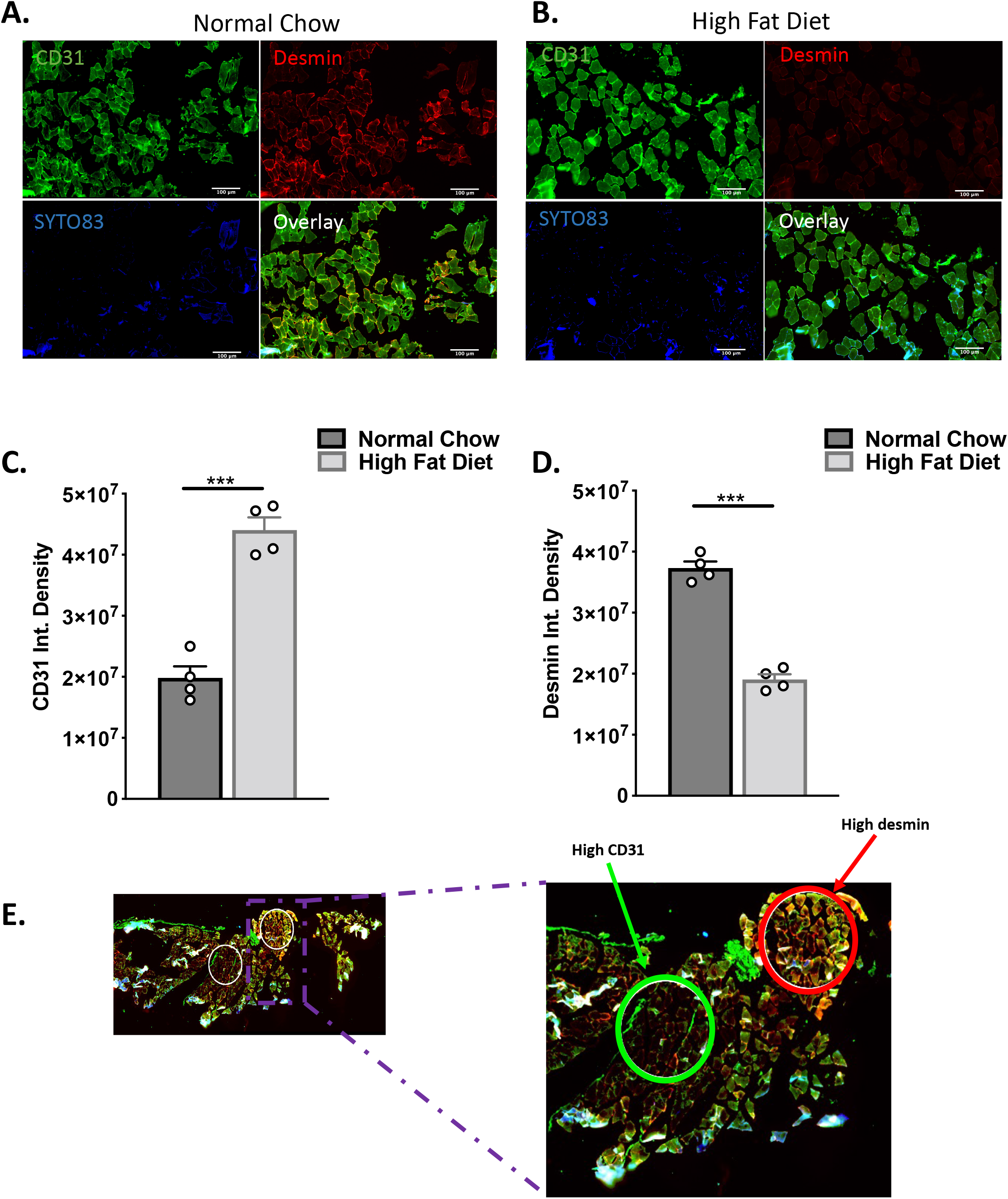
Heterogeneity of murine muscle response to high fat diet. (A-B) Muscle tissue from normal chow and high fat diet mice was subject to immunological staining. Desmin, CD31 and SYTO83 used as morphological markers. (C) Quantification of CD31 abundance in tissue sections. (D) Quantification of desmin abundance. (E) Desmin and CD31 used as morphological markers in normal and high fat diet *tibialis anterior* muscle tissue. 400μm diameter geometric segments selected on tissue sections for spatial profiling. Data presented as means ±SEM. *P<0.05, **P<0.01, ***P<0.0001.

### Diet influences localisation of neuronal and muscle markers in skeletal muscle

In quantifying these regions of interest, we identified elevated CD68, Neun, TMEM119 and GFAP in areas of high desmin and CD31. These proteins were all significantly increased in response to high fat diet exposure. CD68 is commonly expressed in cell populations of macrophages, monocytes, endothelial cells and B and T cells of skeletal muscle (Kosmac *et al*., 2018). Protein abundance unchanged in response to dietary intervention are also shown, Myelin basic protein, CD45, CD39 and CD163 (Fig 4B). The responses recorded here by DSP reflect widespread muscle inflammation during obesity (Fink *et al*., 2014).

**Figure 4.**
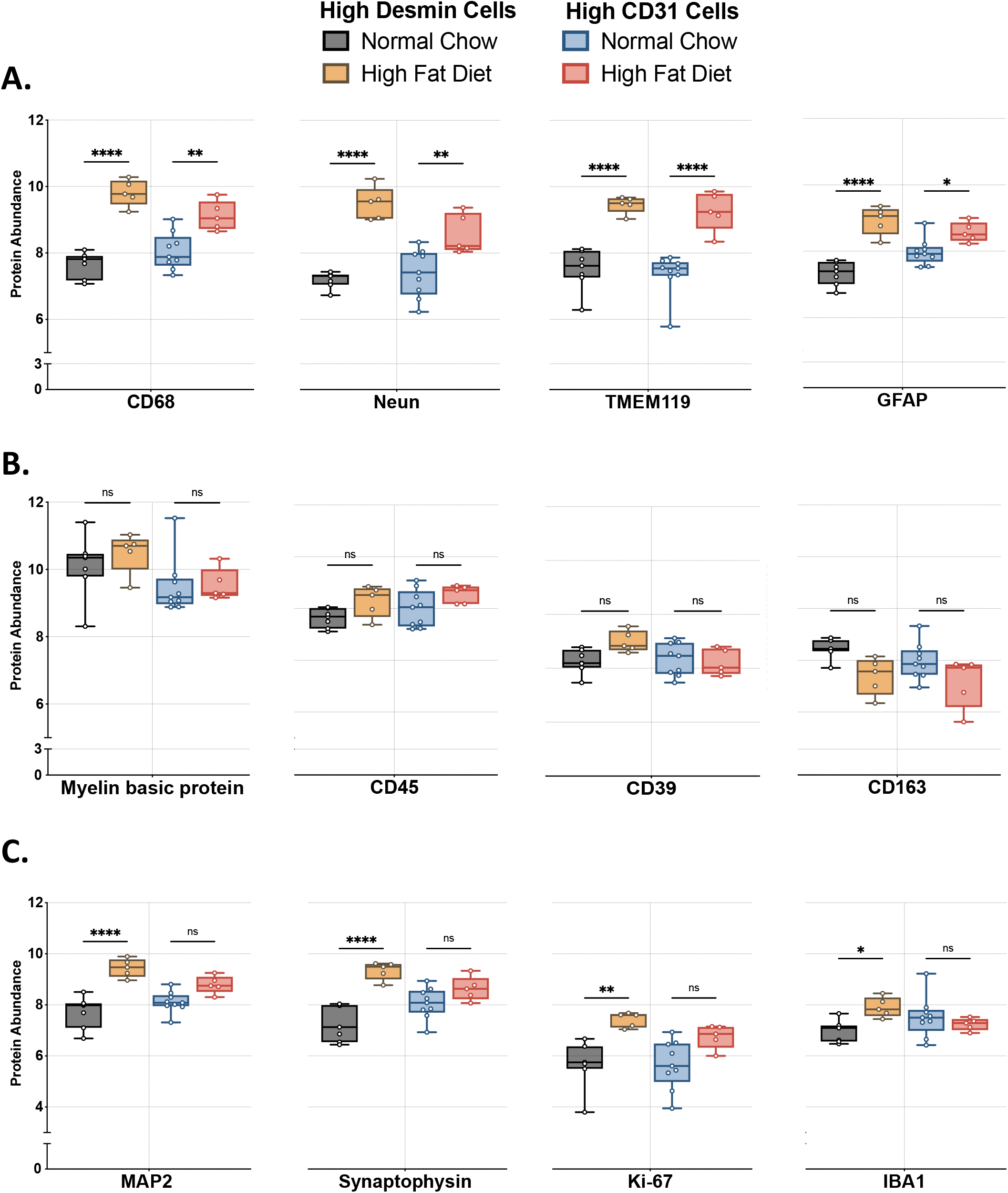
Muscle fibers with elevated CD31 exhibited differential response to high fat diet. (A) Abundance of proteins elevated by high fat diet and independent to spatial position. (B) Proteins unchanged by high fat diet. (C) Proteins elevated by high fat diet in areas of high desmin only. Data presented as box and whisker plots ±SEM. n=5-9 *P<0.05, **P<0.01, ***P<0.0001.

When measuring loci of high desmin it was found that there was significantly elevated expression of MAP2, synaptophysin, Ki-67 and IBA1 (Fig. 4 C). These proteins were all found to have significantly increased abundance in the high fat diet muscle and were spatially resolved to areas of high desmin, not CD31. This indicates that morphologically distinct areas of the response to high fat diet exist within skeletal muscle.

## Discussion

### Spatial changes in neural markers in skeletal muscle tissue exposed to high fat diet

Here we show digital spatial mapping of targeted proteins in murine skeletal muscle. We also document changes in protein abundance in response to high fat diet. We used an established method of dietary induced metabolic stress, through a 60% kcal fat feeding regimen. This generated the typically recorded elevated body weight and circulating blood glucose concentration, reproducing the many studies to have utilised this model (Bryzgalova *et al*., 2008; Ryan *et al*., 2016; Sö Rhede Winzell and Ahré, 2004). Using a panel of markers for neural and inflammatory infiltrates we describe gross changes in spatial abundance of markers including CD31 and MAP2. This supports data demonstrating a dynamic neural and muscle cell-cell relationship, with rapid elevations in angiogenesis in high fat diet fed mice (Silvennoinen *et al*., 2013). Whilst initiating drivers of this are poorly understood, it is evident substantial shifts occur in muscle as a response to dietary intake, and this can be influenced by sex hormones (Martinez-Pena y Valenzuela and Akaaboune, 2020b). These data show skeletal muscle cells may play an important role in plasticity and recruitment of peripheral nervous system structures dictating contractile function. Moreover, a diet high in fat impacts this relationship.

### Areas of elevated CD31 abundance

CD31 is an endothelial cell marker suggesting areas of high abundance are undergoing blood vessel deposition and associated infiltration of neural cells. CD31+ expressing endothelial cells can be recruited to tissue but are also found resident within skeletal muscle (Reyes *et al*., 2006) (Wüst *et al*., 2022). This suggests blood vessel density which is shown to correlate positively with inflammatory infiltration maybe driven in part by muscle residing endothelial cells. Skeletal muscle CD31 is also implicated within muscle cell regeneration suggesting dietary-induced myopathy impacts over satellite cells activation (Huang *et al*., 2018; Motohashi *et al*., 2008; Scalzo *et al*., 2022; Stawerski *et al*., 2013; Yang *et al*., 2021).

### Muscle tissue loci showing increased desmin have shifts in cell markers indicative of neuronal remodelling

Elevated GFAP protein abundance in both higher desmin and higher CD31 geometric segments may imply cellular crosstalk between peripheral glial cells and neurons at the sites of NMJ (Um *et al*., 2017). The same trend was apparent with the postmitotic neuronal marker NeuN, also observed to play a role in muscle plasticity (Kiyoshi Katashima *et al*., 2022). Significantly, increased MAP2 protein, a cytoskeletal neurite marker, was found in the geometric segments of higher desmin in high fat diet fed mice compared to regions of higher CD31. The relationship of desmin expression and heightened neuronal marker requires more attention to determine if a correlation exists in dietary induced-myopathy at NMJ loci.

## Conclusion

Here we detail some of the morphological changes occurring in skeletal muscle in response to a diet of increased fat content. These changes are spatially quantified and reveal increases in regulators of neural cell expression and vascularisation by endothelial activation. These have been further resolved into loci and expression of proteins within distinct areas of CD31 or desmin enrichment. We show areas of high desmin in high fat diet muscle have increase expression of neuronal markers indicative of degradation of neuromuscular junctions and synapse denervation.

With the emergence of spatially resolved protein measures it is anticipated there will be an elevation in the understanding of skeletal muscle turnover and physiology. At present the spatial changes imposed by metabolic stress are poorly understood, with gaps in our appreciation of the molecular drivers and determinants of myopathic states. Quantification by *in situ* of protein abundance will be invaluable in the improved understanding of this dynamic tissue.

## Author contribution statement

LH performed the experimental work. Project was conceived and planed by CLD and RPH. SR and JV aided spatial proteomics. All authors contributed to writing and editing the manuscript.

